# Unearthing modes of climatic adaptation in underground storage organs across Liliales

**DOI:** 10.1101/2021.09.03.458928

**Authors:** Carrie M. Tribble, Michael R. May, Abigail Jackson-Gain, Rosana Zenil-Ferguson, Chelsea D. Specht, Carl J. Rothfels

## Abstract

Testing adaptive hypotheses about how continuous traits evolve in association with developmentally-structured discrete traits, while accounting for the confounding influence of other, hidden, evolutionary forces, remains a challenge in evolutionary biology. For example, geophytes are herbaceous plants—with underground buds—that use underground storage organs (USOs) to survive extended periods of unfavorable conditions. Such plants have evolved multiple times independently across all major vascular plant lineages. Even within closely related lineages, however, geophytes show impressive variation in the morphological modifications and structures (i.e., “types” of USOs) that allow them to survive underground. Despite the developmental and structural complexity of USOs, the prevailing hypothesis is that they represent convergent evolutionary “solutions” to a common ecological problem, though some recent research has drawn this conclusion into question. We extend existing phylogenetic comparative methods to test for links between the hierarchical discrete morphological traits associated with USOs and adaptation to environmental variables, using a phylogeny of 621 species in Liliales. We found that plants with different USO type do not differ in climatic niche more than expected by chance, with the exception of root morphology, where modified roots are associated with lower temperature seasonality. These findings suggest that root tubers may reflect adaptations to different climatic conditions than those represented by other types of USOs. Thus, the tissue type and developmental origin of the USO structure may influence the way it mediates ecological relationships, which draws into question the appropriateness of ascribing broad ecological patterns uniformly across geophytic taxa. This work provides a new framework for testing adaptive hypotheses and for linking ecological patterns across morphologically varying taxa while accounting for developmental (non-independent) relationships in morphological data. [Macroevolution, geophytes, climatic niche evolution, imperfect correspondence]

## Introduction

The evolution of major innovations in life history strategies (how organisms gather and store energy and reproduce) is one of the primary themes of biodiversity research (Adler et al., 2014; Enquist et al., 1999). However, such studies are often limited by methodological challenges in integrating distinct datatypes into a single analytical framework (see Theoretical Overview, below). In this work, we build on recent advances in modeling complex, structured discrete traits and the adaptive evolution of continuous traits to develop an analytical pipeline for testing adaptive hypotheses about the evolution of life history strategies.

In one remarkable example of a life history innovation, certain herbaceous plants can retreat underground by producing the buds of new growth on structures below the soil surface (Raunkiaer et al., 1934), while also storing nutrients to fuel this growth in highly modified, specialized underground storage organs (USOs). Such “geophytes” have evolved independently many times across the plant tree of life, including in diverse and distantly related lineages within the ferns and flowering plants (Tribble et al., 2021). Even within closely related lineages, geophytes show remarkable variation in the particular morphological modifications that allow them to survive underground. By differentially modifying leaves, stems, or roots, geophytes produce complex storage structures through distinct developmental and evolutionary means (reviewed in Tribble et al., 2021). For example, plants may produce bulbs by modifying leaves for storage; rhizomes, corms, and stem tubers by modifying stem tissue; and root tubers by modifying root tissue.

Previous work has suggested that the geophytic habit is correlated with distinct abiotic features—specifically, more seasonal climatic conditions and higher-disturbance regimes (Patterson and Givnish, 2002; Cuéllar-Martínez and Sosa, 2016; Sosa et al., 2016; Sosa and Loera, 2017; Howard et al., 2019)—and may be an adaptation to these conditions (Rees, 1989). Supporting this conclusion, geophytes are particularly diverse in seasonally dry climates such as Mediterranean ecosystems, where they survive hot, dry summers underground and emerge during cool, wet winters to photosynthesize and reproduce, a pattern particularly prominent in the Cape region of South Africa, where almost 15% of native plant species are geophytic (Parsons and Hopper, 2003). Geophytes are also common in deciduous woodland habitats, where their USOs fuel quick spring regrowth to maximize photosynthetic opportunities before trees have leafed-out in spring (Whigham, 2004). One of the first studies to test for a correlation between niche and USO type (Patterson and Givnish, 2002) found that within Liliaceae, plants with bulbs occupy more open and seasonal habitats than plants with rhizomes. In a recent study of monocotyledonous geophytes, Howard et al. (2019) found similar patterns: geophytism is correlated with areas of lower temperature and precipitation and higher temperature variation. Howard et al. (2019) also tested for distinct climate preferences among geophytes with different types of USOs; they found no significant correlations with the exception of an association between rhizomatous geophytes and areas of increased temperature variation. The authors suggest that more detailed morphological data, as well as data related to the developmental origin of USOs (leaf, stem, or root tissue), may be necessary to address variation within geophytes in environmental preferences.

If geophytes with independently evolved and developmentally distinct morphologies converge on the same climatic niche, then the diverse types of USOs may represent different evolutionary paths towards an effectively similar ecological strategy: retreating underground. This result would imply that the diversity of underground forms are due to developmental or genetic constraints that predisposed plants to modify particular types of tissue in different ways when presented with the same types of environmental conditions. Conversely, if plants with different USOs occupy different climatic niches, variation in underground morphology may underlie important differences in how these plants relate to their environment. In this case, geophytism may not be a uniform life-history strategy at all; rather, each distinct morphology may represent a specific adaptive response to a particular set of abiotic conditions. To date, no study has explicitly tested if geophytes with different USOs are adapted to particular climatic niches within an adaptive evolutionary framework that accommodates complex hierarchical relationships among geophyte morphologies.

Representation of diverse geophytic morphologies is particularly high in the monocot order Liliales, which contains roughly 1200 species distributed across the globe (Givnish et al., 2016; Patterson and Givnish, 2002). In this study, we infer a species-level phylogeny of roughly 50% of the taxa in Liliales by capitalizing on the growing availability of published genetic data (Benson et al., 2018), advances in supermatrix construction (de Queiroz and Gatesy, 2007), and model-based tree-building algorithms (Ronquist et al., 2012), which collectively have widened the scope of phylogenetic reconstruction and allow for the inference of increasingly large trees that can provide the statistical power to test complex adaptive scenarios. We use this phylogeny and a newly-developed analysis pipeline to test the relationship between underground morphologies and adaptive evolution to climate seasonality.

### Underground Morphology Background

Geophytic plants produce underground storage organs (USOs) by modifying the main types of vegetative plant tissue: shoots (including leaves and stems) and/or roots. The overall morphology and development of USOs has been reviewed in Tribble et al. (2021); here we summarize the morphologies of taxa in Liliales (those relevant to the present study) and their developmental origins. Geophytes in Liliales may have bulbs, corms, rhizomes, and root tubers. Bulbs are modifications to the shoot system in which leaves serve as the primary storage tissue. These thick, storage leaves form the “layers” visible when a bulb is dissected (such as when cutting an onion). Bulbs are particularly common in Liliaceae (*Lilium*, for example, produce bulbs that survive underground over-winter or during the dry season Rees, 2012).

Corms appear outwardly similar to bulbs but are modifications of the shoot system in which the stem, rather than leaves, serve as the primary storage tissue. Corms are vertically oriented stems with short internodes and uniform swelling along the length of the organ. *Colchicum* (Colchicaceae) grow from corms and are notoriously toxic, especially *C. autumnale*. These corms are often replaced annually by the plant; the nutrients in an old corm are used up as the plant grows a “replacement” corm either apically or laterally (Kubitzki and Huber, 1998).

Other geophytes in Lililales have rhizomes—belowground horizontal stems that may be enlarged (radially) for storage. Rhizomes, like many aboveground stems, have nodes from which branches may emerge, and some rhizomes form extensive belowground networks. The aboveground stems of rhizomatous plants also emerge from nodes along the rhizome. Thus separate aboveground stems may appear to be disconnected individuals from above, but are in actuality linked via the rhizome below (Klimešová et al., 2018).

Some geophytes produce tuberous roots—either enlarged along the length of the root or inflated at the root tip— that also store starch, water, and/or other compounds. In Liliales, these tuberous roots are often produced by plants with rhizomes. Such is the case for most *Bomarea* (Alstroemeriaceae), which frequently grow vines that resprout from underground rhizomes bearing rotund tuberous roots (Sanso and Xifreda, 2001).

### Theoretical Background

While the goal of many phylogenetic comparative methods is to model the evolutionary relationships between (often multiple) traits and species, incorporating diverse data types into a cohesive analytical framework is often stymied by underlying differences in how distinct types of traits are expected to evolve across a tree. Specifically, including continuous and discrete traits—such as climate and morphology—in a single analysis is a longstanding challenge. Some previous approaches to correlate discrete and continuous traits include the use phylogenetic generalized linear models (Garland Jr et al., 1993) and the threshold model (Felsenstein, 2012). However, these methods are purely correlative and do not account for the presence of other, hidden, evolutionary forces that could cause morphological change (Beaulieu et al., 2013; Uyeda et al., 2018).

The Ornstein-Uhlenbeck (OU) process is defined by three parameters (Hansen, 1997; Butler and King, 2004): *σ*^2^ (rate), *θ* (optimum), and *α* (strength of pull). As with Brownian Motion (BM), a continuous trait—such as climate— evolving under the OU process experiences random changes with mean zero and a magnitude proportional to the rate parameter, *σ*^2^. However, unlike BM, a trait evolving under OU is also subject to deterministic changes: it is “pulled” toward the optimal value *θ*, approximating the evolution of the trait toward an adaptive peak. The strength of the pull is proportional to *α* and the distance from the optimum, such that traits that are far away from the optimum are pulled more strongly (the so-called “rubber-band” effect). Further elaboration of the OU process may allow the parameters to vary across the branches of a phylogenetic tree (Beaulieu et al., 2012; Uyeda and Harmon, 2014). In these models, optima may themselves evolve across the phylogeny (Uyeda and Harmon, 2014), unlinked to any observed discrete character. Alternatively, users may specify the location of the multiple optima across the tree (Beaulieu et al., 2012), perhaps based on estimated ancestral states. For example, one could estimate the ancestral states of USO morphology in Liliales and use those estimations to specify the locations of climate regimes across the tree. However, this type of explicit link between trait(s) and adaptive optima attributes all variation in the optima of the continuous trait (*e*.*g*. climate) to the discrete trait (*e*.*g*. USO morphology) and may lead to a spurious correlation. It is therefore necessary to incorporate additional sources of adaptive-optima evolution or to allow for imperfect correspondence between the discrete trait and continuous regimes.

Furthermore, some types of USOs may share greater affinities in morphological and developmental space because they are modifications of the same type of tissue. For example, in both corms and rhizomes, stem tissue is modified for storage, while in bulbs, the primary storage tissue is derived from modified leaves (see Figure 1). A model that represents transitions between corms and rhizomes in the same way that it represents transitions between rhizomes and bulbs effectively erases the complexity of these morphologies and ignores the role that shared developmental mechanisms may play in morphological disparification. Tarasov et al. (2019) presented a novel pipeline—PARAMO— to incorporate developmental hierarchies into discrete ancestral-state estimation using hidden states and structured Markov models (See also Tarasov, 2019). In PARAMO, character states are expanded or combined and hierarchical relationships are expressed by disallowing transitions between certain combinations of character states and requiring transitions through intermediate character states (i.e., using structured, hidden Markov models). Often, these hierarchies are based on ontological definitions, though similar information on character state structure can be incorporated without formal ontologies. This approach addresses the red tail/blue tail problem (Lee and Bryant, 1999): how does one code tail color for a species with no tail? Thus, the PARAMO pipeline is appropriate for modeling the evolution of complex morphologies, where some species may have modifications to certain tissues/ body parts while others lack those modifications or tissues altogether. As such, PARAMO provides a coherent way to model changes in morphologically distinct USOs, such that the type of tissue modified to produce the USOs influences what transitions between structures are allowed. For example, PARAMO models the evolution of rotund root tubers independently from the evolution of rhizomes, as these structures are modifications of different parts of the plants (root and stem, respectively).

**Figure 1:**
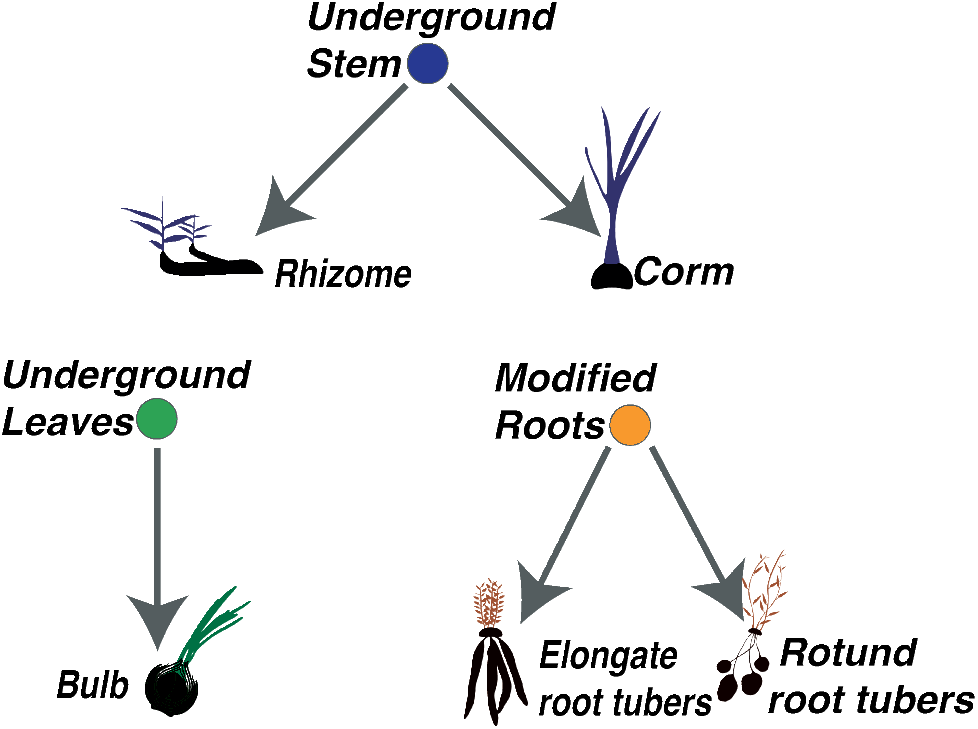
Hierarchy of morphological states used in PARAMO analysis.

Below, we describe our approach to testing the effect of a discrete, morphological trait on the adaptive evolution of a continuous trait while accounting for the nuances presented above.

## Materials and Methods

To test if plants with the same type of USO are evolving towards a shared optimal climatic niche, we developed an analytical pipeline based on recent advances in trait evolution models. We independently model the evolution of discrete and continuous traits. For the discrete trait (USOs), we use a model of morphological evolution that accounts for complex nested relationships among characters (PARAMO, Tarasov et al., 2019). For the continuous trait (climatic niche), we use comparative methods that allow for explicit testing of adaptive hypotheses (bayou; Uyeda and Harmon, 2014). Both these methods produced histories (maps) of trait evolution over a phylogeny, so we then combine these maps to test for correspondence between the traits’ evolutionary histories. Finally, we compare this combined history with one produced under a null model (where there is no relationship between USOs and climate). This procedure allows us to test for correlations between USO evolution and climatic niche evolution while accounting for hiercharchical structure in morphological data produced by developmental processes, the complex adaptive landscape of climatic niche evolution, and the potential effects of hidden evolutionary forces influencing continuous trait evolution.

### Data

We generated three data sets for downstream analysis: a distribution of species-level phylogenies including 50% of the species in Liliales; a modeled climatic niche for each species based on 19 climatic variables (Fick and Hijmans, 2017); and a detailed underground morphology database for all species.

### Phylogeny

We used SUMAC 2.0 (Freyman, 2015) to download gene regions from NCBI GenBank (Benson et al., 2018) for all species in the order Liliales. We targeted genes that clustered with specific guide sequences to obtain sequences for 10 commonly sequenced genes in Liliales (Table S1). We filtered the resulting sequences using custom Python scripts to remove regions that were recovered from fewer than 150 taxa out of the 621 taxa available on GenBank, to remove sites with more than 95% missing data, and to align sequences. All regions were aligned using MAFFT v7.271 (Katoh and Standley, 2013); some alignments (ITS, psbA, and rpl16, and trnL-trnF spacer) failed to align well under MAFFT and were subsequently aligned using PASTA (Mirarab et al., 2015) to improve alignment accuracy. We concatenated the filtered and edited alignments using Sequence Matrix (Vaidya et al., 2011). We reconstructed the phylogeny using MrBayes v3.2.6 (Ronquist et al., 2012) on CIPRES (Miller et al., 2011) with two independent runs of four chains, partitioned by gene region, each under the GTR + Γ model with default priors (Table SS2). We constrained tree space to enforce family-level monophyly and monophyly of the non-parasitic clade (all Liliales families except Campynemataceae and Corsiaceae) according to the Angiosperm Phylogeny Website (Stevens et al., 2016), to reduce run times.

To account for phylogenetic uncertainty, we performed all downstream analyses on a random set of ten trees from this posterior distribution (see section *Sensitivity of Results to Tree Selection*). We dated each of the selected trees in `R` (R Core Team, 2013) using the `chronos()` function from the `ape` package (Paradis and Schliep, 2019)—an implementation of the penalized-likelihood approach (Sanderson, 2002)—using data from two fossils and a secondary calibration (see Table S3; Iles et al. 2015; Givnish et al. 2016). We set *λ* = 1 (the smoothing parameter).

### Climate

We modeled the climatic niche of each sampled species using a newly developed R pipeline, Climate and Niche Distribution Inference (`CaNDI`, available at https://github.com/abbyj-g/candi) that gathers and cleans species occurrences, downloads climate data, and estimates niches for hundreds of taxa at a time. `CaNDI` takes as input a list of species, queries the Global Biodiversity Information Facility (GBIF; Flemons et al., 2007) and the Botanical Information and Ecology Network (BIEN Maitner et al., 2018) for occurrence records, and cleans those records using a series of filters designed to remove latitude and longitude records that fall outside of the species’ native range, are exactly at 0*°*, 90*°*, or 180*°*, or are in the ocean. `CaNDI` retains species with ≥ 5 occurrence points. `CaNDI` then passes these occurrence points and their associated climate data to MaxEnt (Phillips and Dudík, 2008) to estimate the climatic niche. For each species, `CaNDI` returns the probability of occurrence across the landscape.

For the climate data we used 19 bioclimatic variables from the WorldClim database (Fick and Hijmans, 2017), which describe various aspects of temperature and precipitation. Collinearity of predictor variables does not affect model performance (except in cases of model transfer; Feng et al., 2019), so we included all 19 variables in niche estimation. From the niche reconstructions we calculated a single estimate of the optimal value for each climate variable by selecting the value that corresponded to the part of the species’ range with the highest probability of occurrence. Downstream analyses focused on the two axes of the multidimensional niche that describe seasonality: seasonality of precipitation and seasonality of temperature.

### Morphology

We used morphological data from Kew’s World Checklist of Selected Plant Families (WCSP; WCSP, 2020) to describe the USOs associated with each sampled species. For taxa listed as tuberous, we referred to morphological literature (Kubitzki and Huber, 1998; Sanso and Xifreda, 2001; Pate and Dixon, 1982) for more detailed descriptions, as the WCSP uses tuber as a catch-all category, encompassing corms, root tubers, and other organs. The final coding scheme consisted of the presence and absence of eight characters, grouped into three hierarchical clusters based on storage tissue type (leaf, stem, and root; see Figure 1).

### Analysis

We integrated the climate data, morphology data, and phylogeny in a novel analysis pipeline that integrates newly-developed methods for modeling continuous characters (such as climate) and discrete characters (such as morphological categories). This pipeline models the continuous and discrete variables independently, and then asks if variation in adaptive optima for continuous characters are explained by the state of the discrete trait, allowing for complex models of the discrete trait and for an imperfect correspondence between adaptive optima and the discrete trait.

### Climatic Niche Evolution

We used the `R` package `bayou` (Uyeda and Harmon, 2014) to describe the modes of evolution of climatic niche in Liliales across each of ten phylogenies sampled from the posterior of our phylogenetic analysis; `bayou` models the evolution of a continuous character under an OU process (Butler and King, 2004), allowing for the optimum of the OU process to vary across the branches of the phylogeny (Uyeda and Harmon, 2014). Specifically, this approach models the number and placement of adaptive regimes across the branches of the phylogeny, where each adaptive regime is characterized by a unique optimal continuous-trait value, *θ*, rate of evolution, *σ*^2^, and strength of selection, *α*. These regimes and their associated parameter values are sampled in proportion to their posterior probabilities using reversible-jump MCMC (Green, 1995).

For both temperature and precipitation seasonality we log-transformed the variable and ran `bayou` for 3.5 million generations using the priors specified in Table S4. Priors were selected as recommended in the `bayou` tutorial (Uyeda, 2019) except for the prior on *k* (number of *θ*s), which we modified to reflect our uncertainty in *k*. The recommended prior on *k* is a Poisson distribution with *λ* = 10. However, we opted for a geometric distribution with *p*=1/30, which is equivalent to setting an exponential hyperprior on *λ* and increasing the expected value of *λ* from 10 to 30. We assessed convergence using the `R` package `coda` (Plummer et al., 2006) and based on those results discarded the first 1% of samples as burnin. We drew (without replacement) 1000 samples (maps) from the posterior distribution of adaptive regimes for each climatic niche variable to use in subsequent calculations. Each of these samples contains a possible history of adaptive-optima evolution, which maps the adaptive optima along branches of the phylogeny.

### Morphological Evolution

We used the PARAMO pipeline (Tarasov et al., 2019) to reconstruct the evolution of underground morphologies using hidden, structured Markov models and stochastic mapping across the ten trees used in our `bayou` analyses. Underground morphologies are not adequately represented in published ontologies (Tribble et al., 2021; Howard et al., 2021), so instead of using ontologies to determine the hierarchical relationships between states, we built a dependency matrix of USOs based on the type of tissue modified for storage (Figure 1). PARAMO uses hidden states that represent the “predisposition” to evolve USOs of different tissue types (Beaulieu et al., 2013), incorporating the hierarchy of states illustrated in Figure 1. Thus the morphological matrix includes multiple states that correspond to the same observed morphology but different unobserved (hidden) variables. Each cluster (leaf, stem, and root) corresponds to a distinct evolutionary model such that the clusters evolve independently. First, for each of the three clusters, we estimated evolutionary transition rates between character states using the `R` package `corHMM` (Beaulieu et al., 2013). We then reconstructed their evolutionary histories by simulating the evolution of each cluster under the inferred model of evolution and conditioning on the observed data at the tips (i.e., stochastic mapping, Nielsen, 2002; Huelsenbeck et al., 2003) using the `make.simmap()` function in `phytools` (Revell, 2012). Each morphological stochastic map represents one possible scenario of character evolution under the inferred model. We then made a single “combined phenotype” morphological map by overlaying the maps of each cluster. The combined phenotype map illustrates the history of the entire underground phenotype with many character states. We repeated this process 1000 times to produce 1000 combined phenotype maps to capture uncertainty in the history of each cluster.

### Calculating State-Specific Climatic Optima

The previous steps have produced *N* `bayou` maps and *N* morphological maps (where *N* = 1000 in this study). Each morphological map defines a set of breakpoints where the state changes; likewise, each `bayou` map defines a set of breakpoints where the climatic optimum changes. We use the *i*^th^ `bayou` map and the *i*^th^ morphological map to produce a state-specific-optimum map, such that each branch on the tree can be represented as a set of segments and where each segment corresponds to a particular combination of morphological state and climatic optimum (Figure 2B). For a given state-specific-optimum map, we compute the average climatic optimum for a given morphological state *j* as:

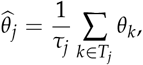

**Figure 2:**
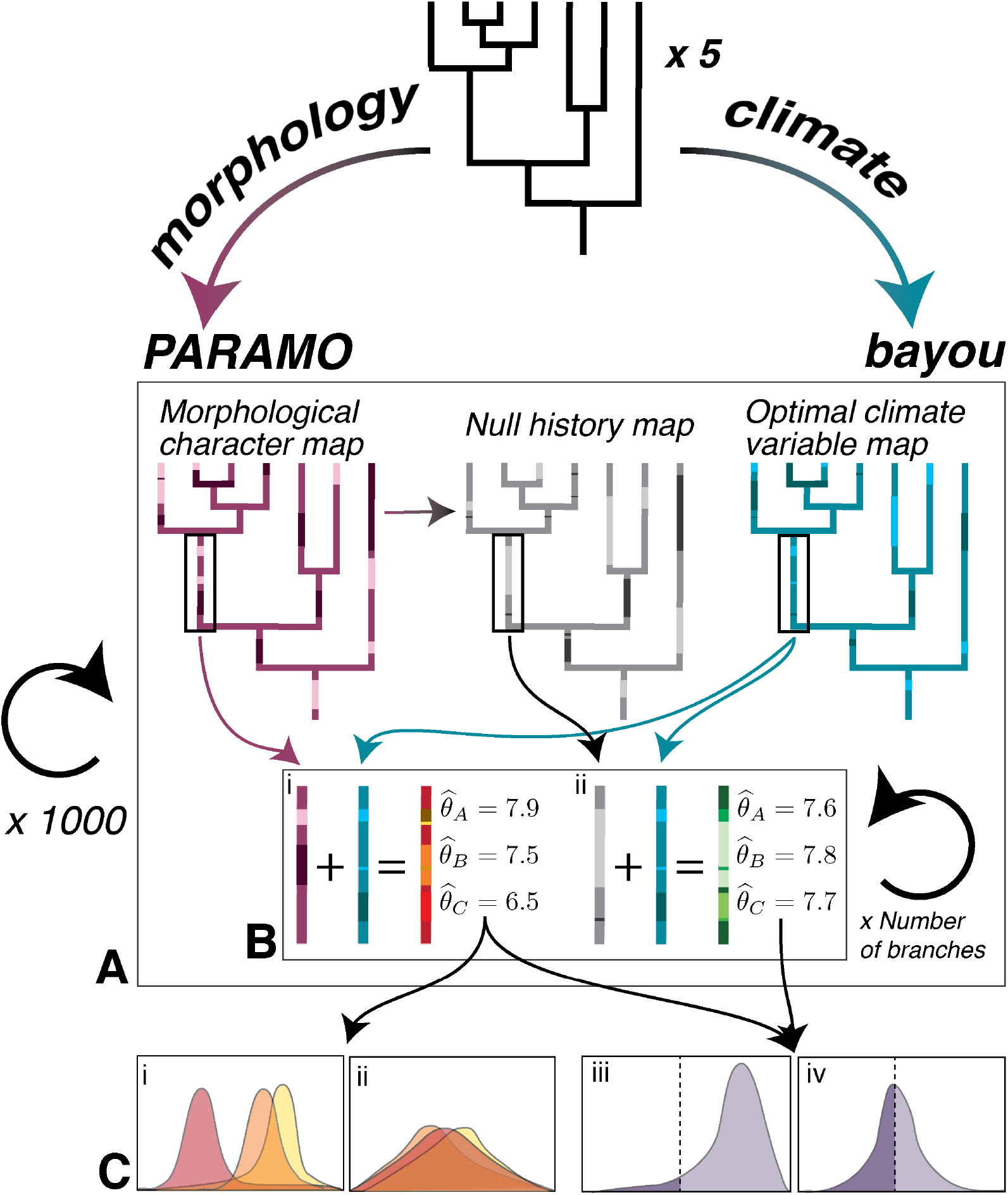
Schematic representation of methods used in this study. For each of ten phylogenies sampled from the posterior distribution, we: A) used PARAMO and bayou to generate 1000 stochastic maps of the morphology and estimated climatic niche optima, individually, and to simulate 1000 null histories (see Methods: Hypothesis testing) using the estimated evolutionary models from PARAMO. B) We combined each of the niche optima maps with a morphological character map, and separately with a null history map, to create two sets of state-specific-optimum maps that show the state-specific 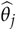 parameters per character state. C) We plot the densities of all estimated state-specific 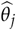 parameters (i, ii) and the distribution of a test statistic (iii, iv; see Methods: Hypothesis testing). Plots i and iii represent a case where we would reject the null model, while in plots ii and iv we would fail to reject the null.

where *T*_*j*_ is the set of all segments in morphological state *j, τ*_*j*_ is the sum of lengths of segments in *T*_*j*_, and *θ*_*k*_ is the climatic optimum associated with segment *k*. We repeat this procedure for each of the *N* state-specific-optimum maps, producing a posterior distribution of state-specific average climatic optima, 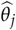.

These statistics represent the relationship between a particular morphology and climate variable. However, our model of morphological evolution (PARAMO) includes multiple states that correspond to the same observed morphology but different hidden variables. As the observed morphology rather than the hidden variable is expected to affect an organism’s relationship to its environment, we combine the state-dependent optima of states with the same observed morphology but different hidden variables before computing the above statistics.

Some states are visited infrequently during stochastic mapping and thus have high percentages of missing data (*i*.*e*., were rarely inferred as ancestral states across the 1000 maps), which makes estimates of their state-specific optima more uncertain. To avoid these uncertain estimates, we do not compute the state-specific optima statistics for states represented in less than 50% of the maps.

### Hypothesis Testing: Climate and Morphological Trait Correlation

Our null hypothesis is that plants with different USOs are not evolving towards different climatic niche optima; in other words, that state-specific climatic niche optima (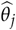) are equal. Our alternative hypothesis is that plants with different USOs have different state-specific optima, 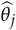. However, for any finite dataset, inferred state-specific 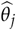 values will be different, even under the null hypothesis, and this issue is exacerbated by the phylogenetic structure of the inferred 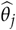 values. To account for this structure, and the finite sample, we calculated a test statistic (defined below) that summarizes the overall difference between any particular set of state-specific 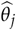. We simulated the distribution of that test statistic under the null model and checked whether the differences generated under the null model were significantly less than the differences of the empirical estimates.

We simulated 1000 null histories of the morphologies using the `sim.history()` function in the `R`package `phytools` (Revell, 2012) and the estimated Q-matrices from the empirical analyses. This procedure differs from stochastic mapping in the empirical analysis in that the simulations are not conditioned on the observed character data. As in the empirical analysis, we also combined the three morphologies to produce 1000 null histories of the combined phenotype. This null model represents the case where a discrete trait evolves under the same evolutionary model as the observed discrete trait, but without the observed pattern at the tips. Any correspondence between the simulated traits and climate is due to chance, the distribution of the climate data on the tree, and/or the rates of evolution and stationary frequencies of the model, rather than the distribution of morphological states across the tree.

To calculate the test statistic, for a given vector of state-specific 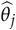 values, we calculate the pairwise distance between all state-specific 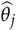. These distances represent the extent to which character states are linked to different estimated 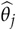; in other words, the optimal climatic values differ between discrete states. These distances result in a distance matrix, *D* (for a simple two-state example):

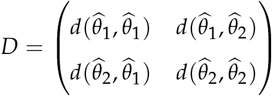

where 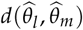 is the distance between 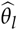 and 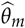:

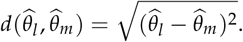

We then measure the overall amount of difference using the Frobenius norm, *F*(*D*), which summarizes the magnitude of the distance matrix. For a given `bayou` map *i*, we calculate *S*_*i*_, the difference between *F*(*D*_*i*_) given the stochastic map and *F*(*D*_*i*_) given the null history. We then compute *S*_*i*_ for each `bayou` sample; if 0 is in the 95% probability interval of *S*, we do not reject the null model.

We performed this hypothesis test for each combination of climate seasonality (precipitation and temperature) and morphology (leaf, stem, and root, as well as the combination of all three), for a total of eight comparisons.

### Sensitivity of Results to Phylogenetic Uncertainty

We performed all the above analyses on ten trees randomly sampled from the posterior distribution of our phylogenetic analysis. To determine if our results were sensitive to particularities of the sampled trees, we analyzed each tree individually and two sets of five trees each. While the results vary by individual tree (Figure 5), sampling at least five trees is sufficient to produce consistent results (Figure S2); the results do not change meaningfully between either analysis of five trees or the full analysis of ten trees (Figures 5 and S2), so we present results based on the combined ten trees.

### Data and Code Availability

All code is freely available on GitHub: https://github.com/cmt2/underground_evo and all data will be made available on Dryad prior to publication.

## Results

Across all ten trees, the ancestor of Liliales is estimated to have had a rhizome with no leaf or root modifications (Figure 3 A–D shows the reconstructions for one tree), in agreement with previous work (Patterson and Givnish, 2002; Howard et al., 2019). For both temperature and precipitation seasonality, the estimated optima at basal branches in the phylogeny are highly seasonal, with subsequent shifts into less seasonal optima along more recent branches (Figure 3 E and F).

**Figure 3:**
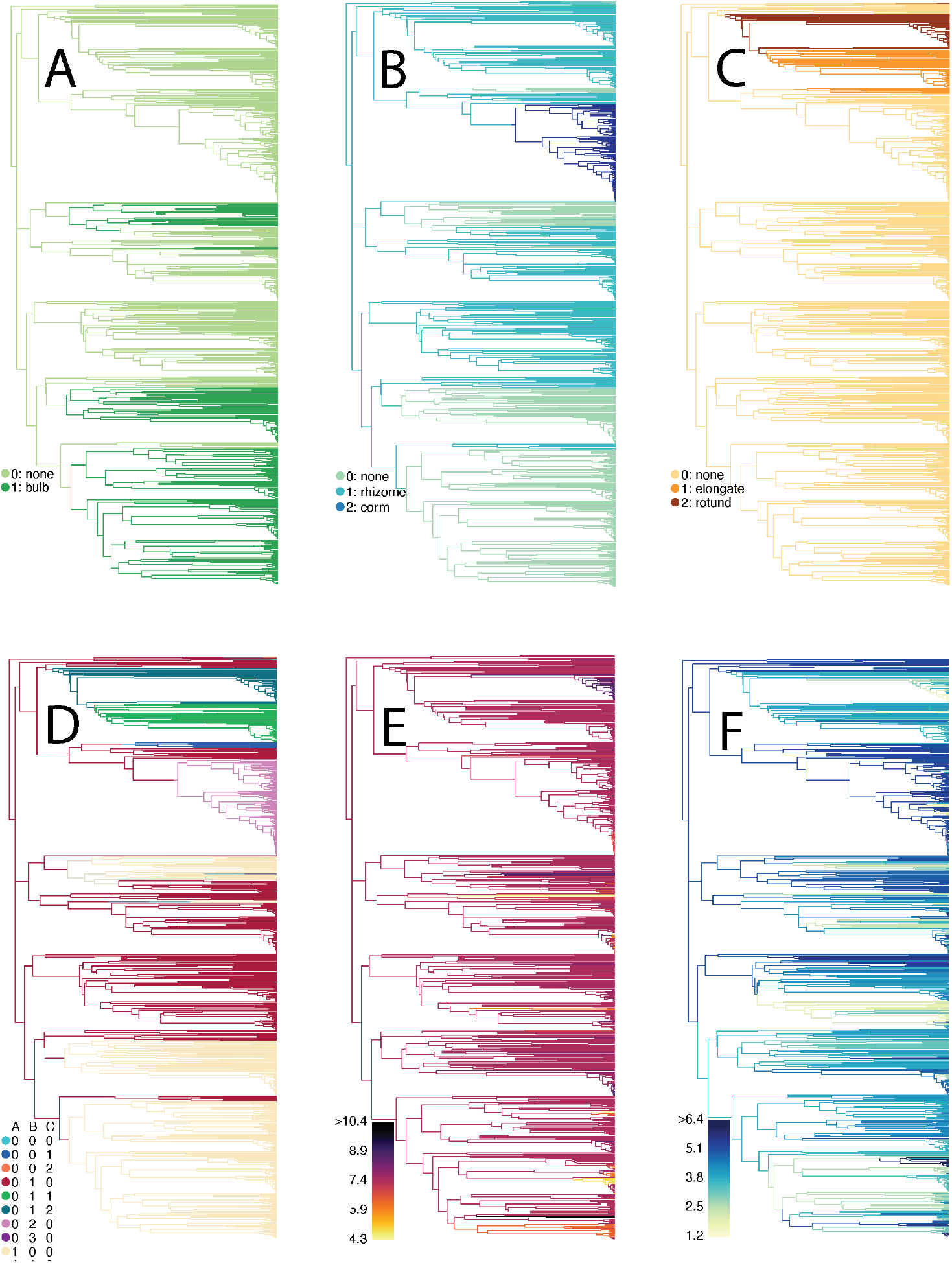
Stochastic maps of estimated ancestral states for one of five trees used in the analysis: (A) leaf character map showing inferred evolutionary history of having or not having a bulb, (B) map of estimated history of stem modifications, (C) map of estimated evolutionary history of root modifications, (D) combined phenotype character produced by amalgamating maps A-C, (E) posterior mean branch-specific *θ* for temperature seasonality, (F) posterior mean branch-specific *θ* for precipitation seasonality. In A, B, and C, the “none” category applies to absence of modifications of the relevant tissue type; those taxa may have modifications of the other tissues. In D, the legend rows correspond to states and the legend columns correspond to the individual characters mapped in A (leaf), B (stem), and C (root) above. Thus 000 refers to none for leaf, stem, and root (the absence of any USO), and 012 refers a geophyte with rhizomes and rotund root tubers.

Figure 4 illustrates the distribution of estimated state-specific optima. Overall, the distributions of state-specific optima are more dispersed across the temperature seasonality axis than the precipitation seasonality axis. For the leaf cluster, the estimated averaged optima for bulb-bearing plants are more seasonal than those for no bulb. For the stem cluster, plants with either rhizomes or corms are estimated to be associated with reduced seasonality than plants without stem modification, though for precipitation only rhizomes are estimated to be less seasonal than the other states. In the root cluster, plants with root tubers (especially rotund root tubers) have lower temperature seasonality than those without root tubers, but for precipitation, plants with rotund root tubers and without root tubers overlap in their distributions while plants with elongate root tubers are estimated to be more seasonal. In the combined phenotype for temperature, the most seasonal state for precipitation is plants with both bulbs and rhizomes and the least seasonal state is plants with rhizomes and rotund root tubers. Plants with bulbs alone, with rhizomes alone, and with both bulbs and rhizomes are the three most temperature seasonal states. For precipitation seasonality in the combined phenotype, most state-specific optima have highly overlapping distributions, though it appears that plants with rhizomes are slightly less seasonal and plants with rhizomes and elongate root tubers are slightly more seasonal than other distributions. For both temperature and precipitation, the distribution corresponding to no modified underground organs (non-geophytes) falls out intermediate along both climatic niche axes and overlaps with many other states, signifying that non-geophytes are not more or less seasonal than geophytes.

**Figure 4:**
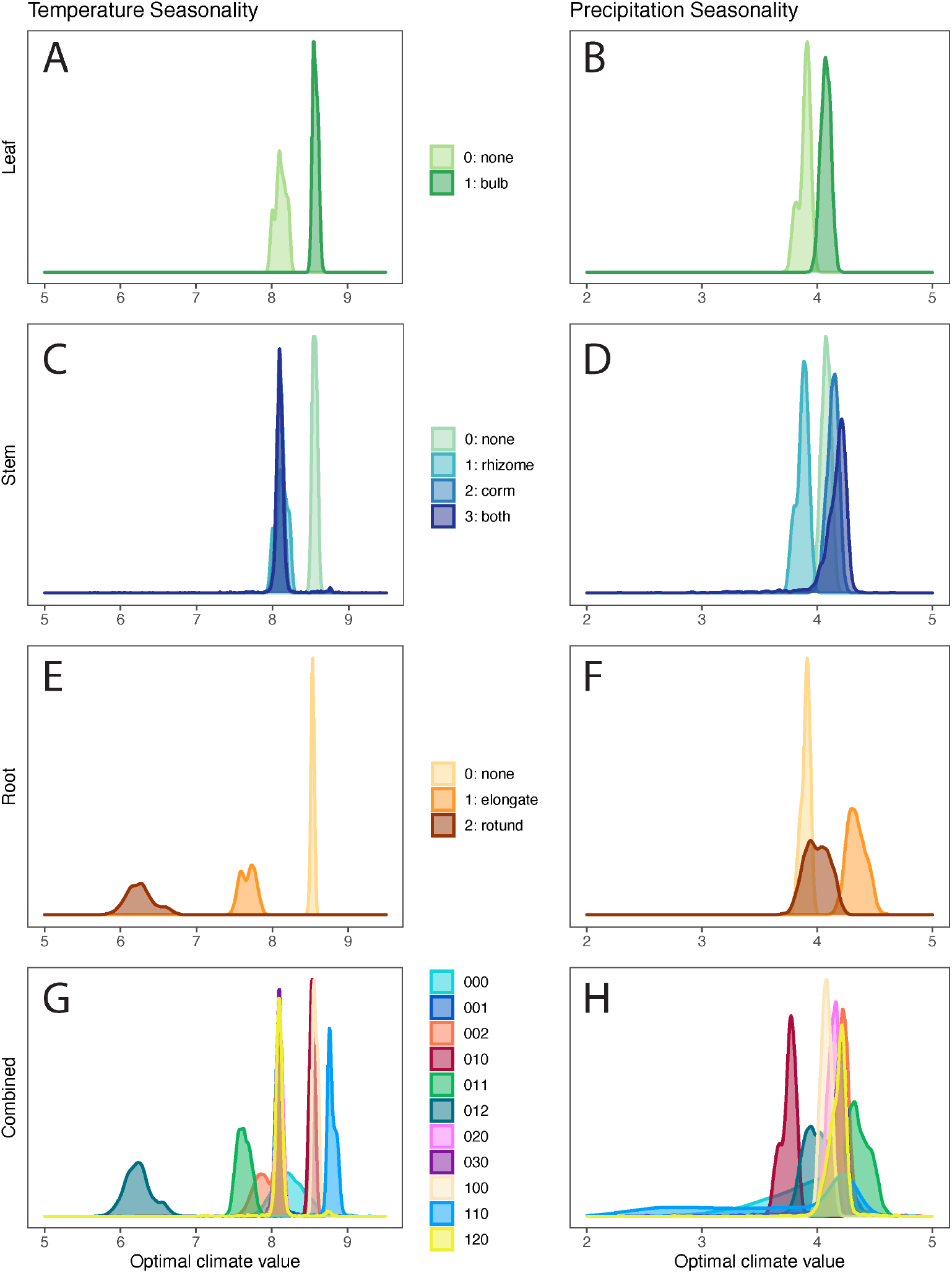
Posterior distributions of state-specific 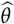 parameters. Panels A, C, E, and G depict 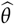 distributions for temperature seasonality, while panels B, D, F, and H show 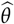 distributions for precipitation seasonality. A — B refer to the leaf cluster, C — D to the stem cluster, E — F to the root cluster, and G — H to the combined phenotype. For these combined phenotype densities (G — H), digits in the state labels refer to states for each cluster. The first digit corresponds to the leaf state, the second digit corresponds to the stem state, and the third digit corresponds to the root state. For example, the 000 state refers to the absence of any USO, and 012 refers to a geophyte with rhizomes and rotund root tubers.

While the state-specific optima curves in Figure 4 appear distinct for many of the morphology-climate comparisons, across both climate variables, no distributions are more different than expected by chance (P-values 0.216–0.626), with the exception of the root cluster for temperature seasonality, which is significant (P-value 0.044; Figure 5).

**Figure 5:**
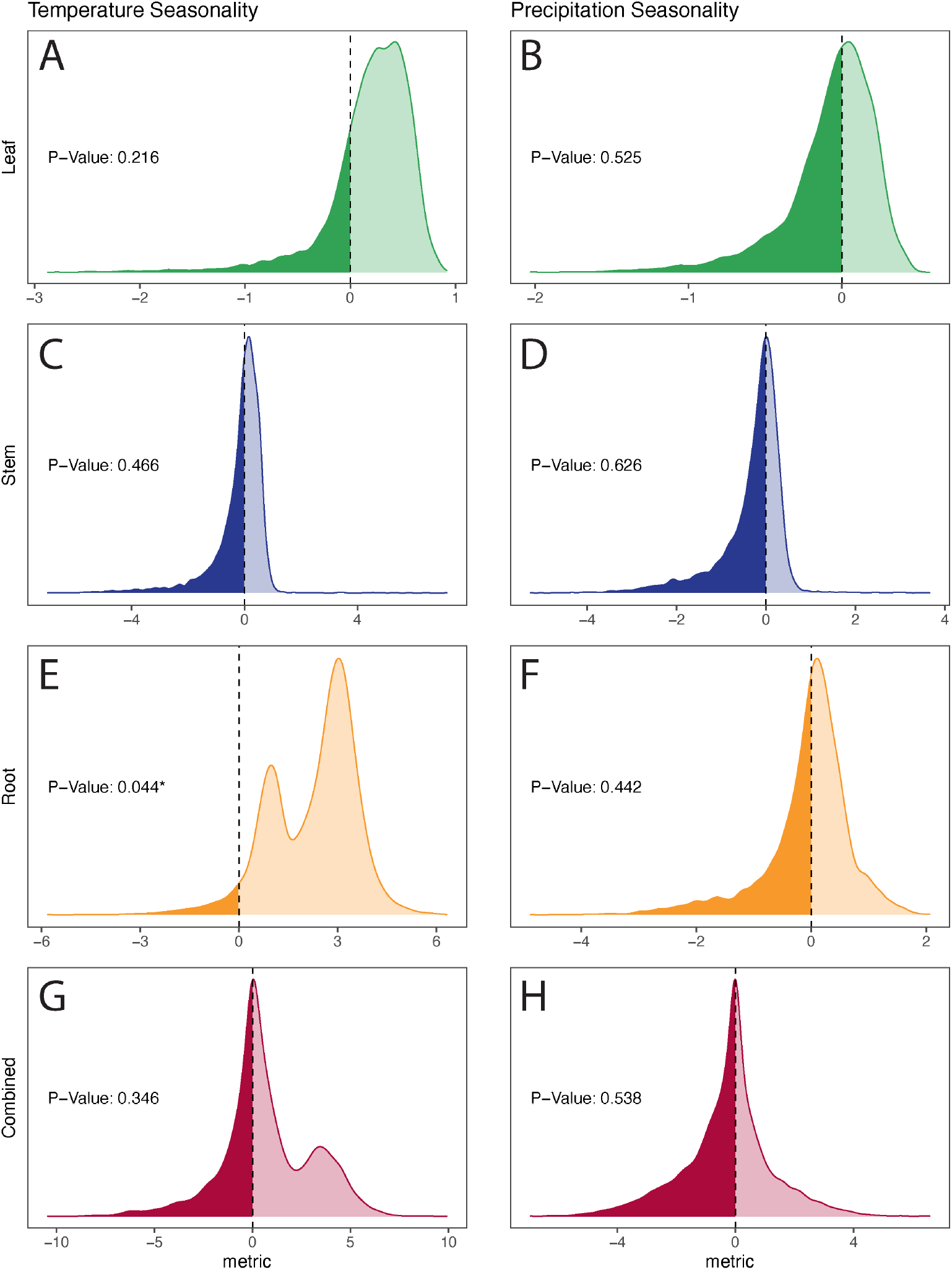
Test statistic (*S*) distributions. Panels A, C, E, and G depict *S* distributions for temperature seasonality, while panels B, D, F, and H show *S* distributions for precipitation seasonality. A — B refer to the leaf cluster, C — D to the stem cluster, E — F to the root cluster, and G — H to the combined phenotype. Distributions with 95% ≥ 0 are considered evidence for a statistically significant difference in state-specific 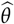. P-Values correspond to the percent of values equal to or less than zero. Of all comparisons, only root and temperature seasonality are statistically significant (panel E).

## Discussion

### Applicability of analysis pipeline for future comparative studies

This work demonstrates the utility of applying complex evolutionary models to address a long-standing challenge in statistical comparative biology: modeling relationships among discrete and continuous traits. Some implementations of OU models allow for continuous traits to evolve under regimes dictated by the evolutionary history of discrete traits (such as implemented in `OUwie`, Beaulieu et al., 2012); an important advancement in these methods allows for modeling a hidden character with a fixed number of states, thus including hidden variation in the evolution of the continuous trait (Vasconcelos et al., 2021). However, these approaches impose potentially restrictive assumptions on the number of adaptive regimes associated with either hidden or observed traits.

Our analytical pipeline unites three main components. First, we use PARAMO (Tarasov et al., 2019) to estimate the evolutionary history of a discrete trait while taking into account hierarchical relationships between character states. Second, we use `bayou` (Uyeda and Harmon, 2014) to estimate the number and location of adaptive regimes of continuous-trait evolution that best fit the data. Third, our pipeline combines these estimates by calculating the average optimal values for each discrete trait and asks if these averaged optima are more different from each other than expected under a null model.

This pipeline has several advantages over existing methods. First, the model parameters have clear biological interpretations. Densities from Figure 4 show the distributions of probable optimal phenotypes (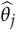) of the continuous trait by discrete category. These 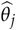 correspond to peaks in the adaptive landscape of the continuous trait, so the estimated parameter values have a clear and direct link to evolutionary theory, unlike the estimated effect sizes from linear models. Second, the method directly models the evolutionary process of the discrete traits and thus is appropriate for addressing evolutionary dependence between continuous and discrete traits. This contrasts with methods such as phylogenetic ANOVA, which treat discrete tip states as purely a source of nonrandom structure at the present, rather than actually modeling the evolution of the trait across the tree. Third, the pipeline can accommodate a wide variety of models for the evolution of the discrete traits. We use PARAMO (Tarasov et al., 2019) to model the nested relationships among USOs derived from the same tissue. In some empirical cases, not including the present study, traits are explicitly defined in ontologies (e.g., phenoscape.org), and PARAMO uses these definitions directly to establish the hierarchy between character states. As the use of ontologies in comparative biology becomes more common, incorporating hierarchies codified in ontologies will greatly expand the ability of researchers to test increasingly complex hypotheses, including how developmental processes impact trait evolution and the relationships among traits (Howard et al., 2021). However, any model of discrete trait evolution that can produce simulated character histories can also generate stochastic map histories, and thus could be used instead of PARAMO. For example, one could produce stochastic maps from a state-dependent speciation and extinction (SSE) analysis (BiSSE, for example, Maddison et al., 2007), and thus link discrete ancestral states, informed by variable speciation and extinction rates, to a continuous character of interest.

Finally, this method allows for imperfect correspondence between the estimated optimal continuous trait and the discrete states. The evolution of continuous traits in general, and climatic niches in particular, is complex and almost certainly explained by many other variables beyond those that are targeted for analysis. It is thus inappropriate to force our model to test between either no correspondence or full correspondence between the continuous-trait evolution and the state of a focal discrete trait. SSE methods may provide a useful analogy; the addition of hidden states allow these models to apportion the variation in speciation and extinction rates between the observed trait of interest and an unobserved hidden trait (Rabosky and Goldberg, 2015; Beaulieu and O’Meara, 2016). Similarly, our method examines the distributions of state-dependent optima without requiring that the continuous trait regimes and the discrete trait vary perfectly together over the tree. We test for significance by comparing these observed distributions to distributions that are built from our null expectation of random evolution of the discrete trait. We thus avoid the “straw-man effect” described in May and Moore (2020), in which any variation in the process of continuous-trait evolution is spuriously attributed to the discrete trait, because the null hypothesis is that there is no variation at all.

As with all current approaches to modeling the correlated evolution of traits along phylogenies, this method may be susceptible to false positives caused by phylogenetic “psuedoreplication”. If both of the two traits of interest happen to transition between states one time, at the same point on the phylogeny, the stochastic maps will match perfectly. Our approach will likely infer a strong degree of correspondence between the traits (though the exact degree of correspondence will depend on the null model). However, if each trait has undergone singular shifts, it is possible that this association is either driven by a third, unobserved trait, or by chance alone. This “psuedoreplication” is an active area of research (see Maddison and FitzJohn, 2015; Uyeda et al., 2018); in their review of this issue, Uyeda et al. (2018) advocate for careful examination of datasets and emphasize the importance of examining patterns of trait evolution of individual characters when testing for correlated evolution. Importantly, our analytical pipeline facilitates this examination by producing stochastic maps of the discrete character and continuous character regimes evolving across the tree, so that singular associated shifts will be straightforward to identify.

All code and scripts used in this pipeline are publicly available at https://github.com/cmt2/underground_evo.

### Getting at the root of the problem: are USOs adaptations to particular climatic niches?

We used this pipeline to test for correlated evolution between climatic seasonality and geophytic USOs. These analyses demonstrate that plants in Liliales with the same underground storage organ do not occupy different climatic niches more than expected by chance, with the exception of root morphology, where the presence of modified roots, especially rotund root tubers, is associated with lower temperature seasonality. Furthermore, non-geophytes in Liliales are not associated with more or less seasonal climates than their geophytic counterparts. While many of the state-specific optima curves appear distinct (Figure 4), the null model suggests that these differences could be observed even with no correspondence between the trait and climate (Figure 5), due to the phylogenetic structure of environmental niche preference in the underlying data.

These findings suggest that root tubers may be an adaptation to distinct ecological conditions or that root-tuber-bearing taxa experience physiological constraints that restrict them to particular climatic niches, unlike the other geophytes included in this study. Root tubers are also unique morphologically among the USOs included in this study; unlike corms, rhizomes, and bulbs, root tubers—especially in Liliales—are rarely the source of perennating underground buds. Most geophytes in Liliales with root tubers also have rhizomes (e.g., *Bomarea, Alstroemeria*), which may serve as the source of underground buds while the root tubers store nutrients and water (Tribble et al., 2021). Many geophytes regenerate their USOs annually (especially bulbs and corms; Pate and Dixon, 1982; Kamenetsky and Okubo, 2012), and thus the processes of nutrient flow between the USO and the above-ground plant are necessarily linked to the same seasonal cycles. Partitioning growth and storage between organs (as in the case of the species with tubers and rhizomes) may be particularly advantageous in climates with less temperature seasonality, as it allows for the continuous production of aerial shoots and the periodic replacement of stored nutrients as needed. Alternatively, places with less seasonal temperatures may be less likely to reach the freezing point, and root tubers may be particularly maladapted to frost compared to USOs derived from stem or leaf tissue.

The results of our study differ from previous work in three primary ways. First, while we found no significant difference in the environmental niche of geophytes and non-geophytes, prior work suggested that geophytes are associated with lower temperatures and precipitation and higher temperature variation compared with non-geophytes (Howard et al., 2019). Because of the sparse representation of non-geophytes in Liliales, our dataset may lack sufficient power to detect generalizable patterns of climatic niche occupancy between geophytes and non-geophytes. Secondly, previous work found that rhizomes are correlated with increased temperature variation and found no evidence for different niches between tuberous and non-tuberous taxa (Howard et al., 2019), while our analysis found no significant association between rhizomatous or non-rhizomatous taxa but instead suggests that taxa with root tubers evolve towards lower optimal values of temperature seasonality. These differences may be due to the scale of the study; Howard et al. (2019) aimed to identify monocot-wide patterns, but our study focuses a smaller taxonomic scale, and is able to investigate patterns of morphological evolution in greater detail. Of particular note, the Howard et al. (2019) study does not distinguish between root tubers and stem tubers, and thus would not have been able to recover the association between roots and lower temperature seasonality that we find here. Thus, our results illustrate the importance of detailed “development-aware” character coding in comparative studies.

Thirdly, Patterson and Givnish (2002) found evidence that in the core Liliales, convergence on bulbs correlated with independent transitions into seasonal and high-light habitats, but while we also find evidence for several independent transitions to bulbs, the association between bulbs and increased seasonality is not statistically significant in our results (P-values 0.216 and 0.525 for temperature seasonality and precipitation seasonality respectively). The Patterson and Givnish (2002) study was one of the first to address geophyte evolution in a phylogenetic framework and correlated storage organs with discrete habitat types, but recent advances in statistical phylogenetics have made more nuanced approaches possible, such as our new pipeline. Specifically, our study uses continuous climatic data directly in the analysis, rather than discrete habitat categories, and employs a coherent model of adaptive continuous trait evolution. It is possible that discrete habitat categories obscured important variation in seasonality, leading to a spurious correlation between seasonality and the presence of bulbs.

In Liliales, root tubers are mostly restricted to a few clades (namely *Bomarea, Alstroemeria, Burchardia*, and a few additional taxa), so it is possible that the strong association between decreased temperature seasonality and the presence of root tubers is due to “psuedoreplication” and is driven by an unmeasured trait that happens to co-occur in those clades (see discussion of “psuedoreplication” above). For this reason, extrapolating our results to non-liliid geophytes may not be appropriate, and follow-up studies should address variation in root morphology and climate in other clades, particularly in groups with many independent transitions between the absence and presence of root tubers and between different root tuber morphologies. *Asparagus* would be a particular appropriate system in which to further test these associations, as root morphology is highly variable in the genus (Leebens-Mack, pers. comms.). However, there are few other clades known for well-characterized variation in the presence and absence of root tubers or for variation in root tuber morphology. Underground morphology is vastly under-characterized in many plant clades (Tribble et al., 2021; Janzen et al., 1975), so this work also motivates increased morphological characterization of USOs and root morphology in particular, and demonstrates the importance of continued emphasis on classic botanical techniques for understanding biodiversity. In addition, studies that characterize the functional ecology and physiology of root tubers will likely yield important insights into how and why they differ from other USOs.

## Conclusions

This study introduces an analysis pipeline that infers the relationship between adaptive optima for a continuous trait and the hierarchical, nested evolution of a discrete trait, controlling for other factors driving changes in adaptive optima. While previous methods have attempted to address this issue, ours it the first that allows the number of adaptive regimes to vary in a way that best fits the data. This pipeline is applicable across many areas of evolutionary biology and may serve as a model for future hypothesis-driven comparative research. Our pipeline can accommodate an array of complex models of discrete morphological trait evolution (such as models that account for the developmental history of morphological traits) into tests of correlated trait evolution, and implements a novel approach to account for imperfect correspondence between adaptive regimes and discrete traits. These advances are key steps forward as ecological and evolutionary studies increasingly seek to incorporate the nuance and complexity of natural variation into quantitative models that permit formal hypothesis testing.

## Author Contributions

CMT designed and executed the study. CMT collected data. CMT and AJ-G performed analyses, and MRM advised on statistics. CMT, MRM, CJR, CDS, and RZ-F guided interpretation of results.

## Acknowledgements

We would like to acknowledge Benjamin K. Blackman, David D. Ackerly, members of the Rothfels Lab sensu lato at UC Berkeley, and two anonymous reviewers for valuable feedback on earlier versions of the manuscript. Conversations with Cody Coyotee Howard and Jesus Martínez-Gómez guided treatment of underground storage organs and interpretation of results. Josef C. Uyeda provided advice on implementing PARAMO without relying on formal ontologies. This is publication number xxx from the School of Life Sciences at UHM.

