## Supplemental Materials for "Unearthing modes of climatic adaptation in underground storage organs across Liliales"

### Supplemental Material

#### Phylogenetic reconstruction

**Table S1:** List of gene regions and percent coverage used in phylogenetic reconstruction.

| Gene Region | Genome | Taxon Coverage | Length (bp) |
|---|---|---|---|
| matK | chloroplast | 51.2% | 1699 |
| trnL-trnF spacer | chloroplast | 50.6% | 1169 |
| ITS | nuclear | 35.6% | 979 |
| atpB | chloroplast | 33.7% | 1500 |
| psba | chloroplast | 28.2% | 1127 |
| rpl16 | chloroplast | 26.9% | 1371 |
| rbcL | chloroplast | 21.9% | 733 |
| nadhF | chloroplast | 21.7% | 701 |
| atpB-rbcL spacer | chloroplast | 20.3% | 940 |
| rps16 | chloroplast | 19.8% | 868 |

**Table S2:** Priors for Bayesian phylogenetic analysis in `MrBayes`.

| Parameter | Prior |
|---|---|
| Revmat{n} | Dirichlet(1.00,1.00,1.00,1.00,1.00,1.00) |
| Pi{n} | Dirichlet(1.00,1.00,1.00,1.00) |
| Alpha{n} | Exponential(1.00) |
| Ratemultiplier{all} | Dirichlet(1.00,1.00,1.00,1.00,1.00,1.00,1.00,1.00,1.00,1.00) |
| Tau{all} | Prior on topologies obeys constraints |
| V{all} | Unconstrained:GammaDir(1.0,0.1000,1.0,1.0) |

**Table S3:** Constraints used for dating analysis.

| Constraint | Age (mya) | Calibration Type |
|---|---|---|
| *Luzuriaga* stem node | 23.2 | Fossil from Iles et al. (2015) |
| *Ripogonaceae* stem node | 51−52 | Fossil from Iles et al. (2015) |
| Liliales stem node | 115.6−131.1 | Secondary from Givnish et al. (2016) |

### Climate analysis with `bayou`

**Table S4:** Priors used for `bayou` analysis of climate variables. In this context, climate refers to the particular climatic niche variable used in the analysis, so the priors on $\theta$ differed for the two analyses.

| Model Parameter | Distribution | Distribution Parameter Value(s) |
|---|---|---|
| $\alpha$ | Half Cauchy | $scale = 0.1$ |
| $\sigma^2$ | Half Cauchy | $scale = 0.1$ |
| $K$ | Geometric | $p = 1/30$ |
| $\theta$ | Normal | $mean = mean(climate)$ |
| | | $sd = 1.5 \cdot sd(climate)$ |

### Consistency of results across phylogenies

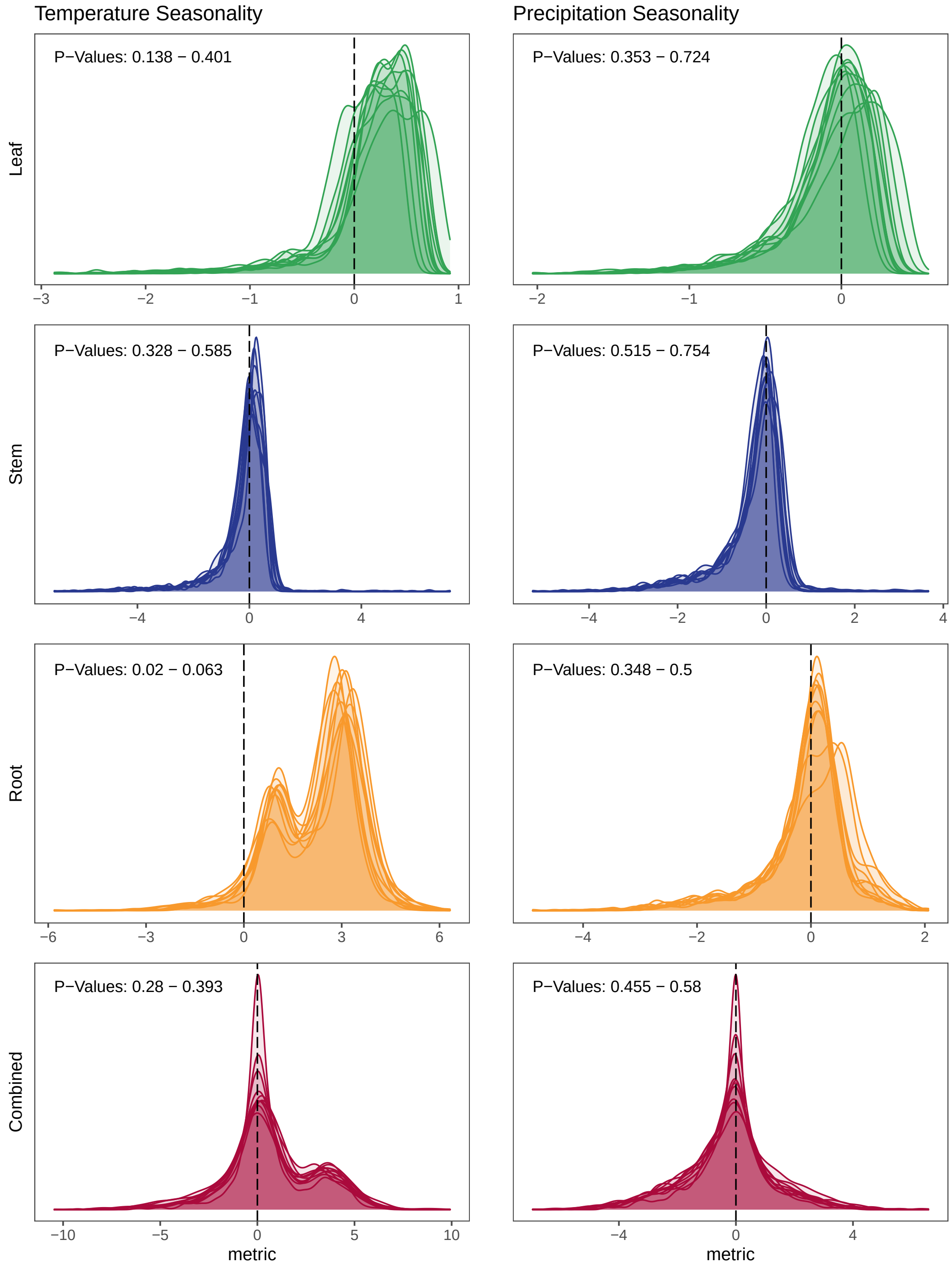


**Figure S1:** Test statistic ($S$) distributions for each of the ten phylogenies in the analysis. Distributions with 95% $>$ 0 are considered evidence for a statistically significant difference in state-specific $\theta$. P-Values correspond to the percent of values equal to or less than zero.

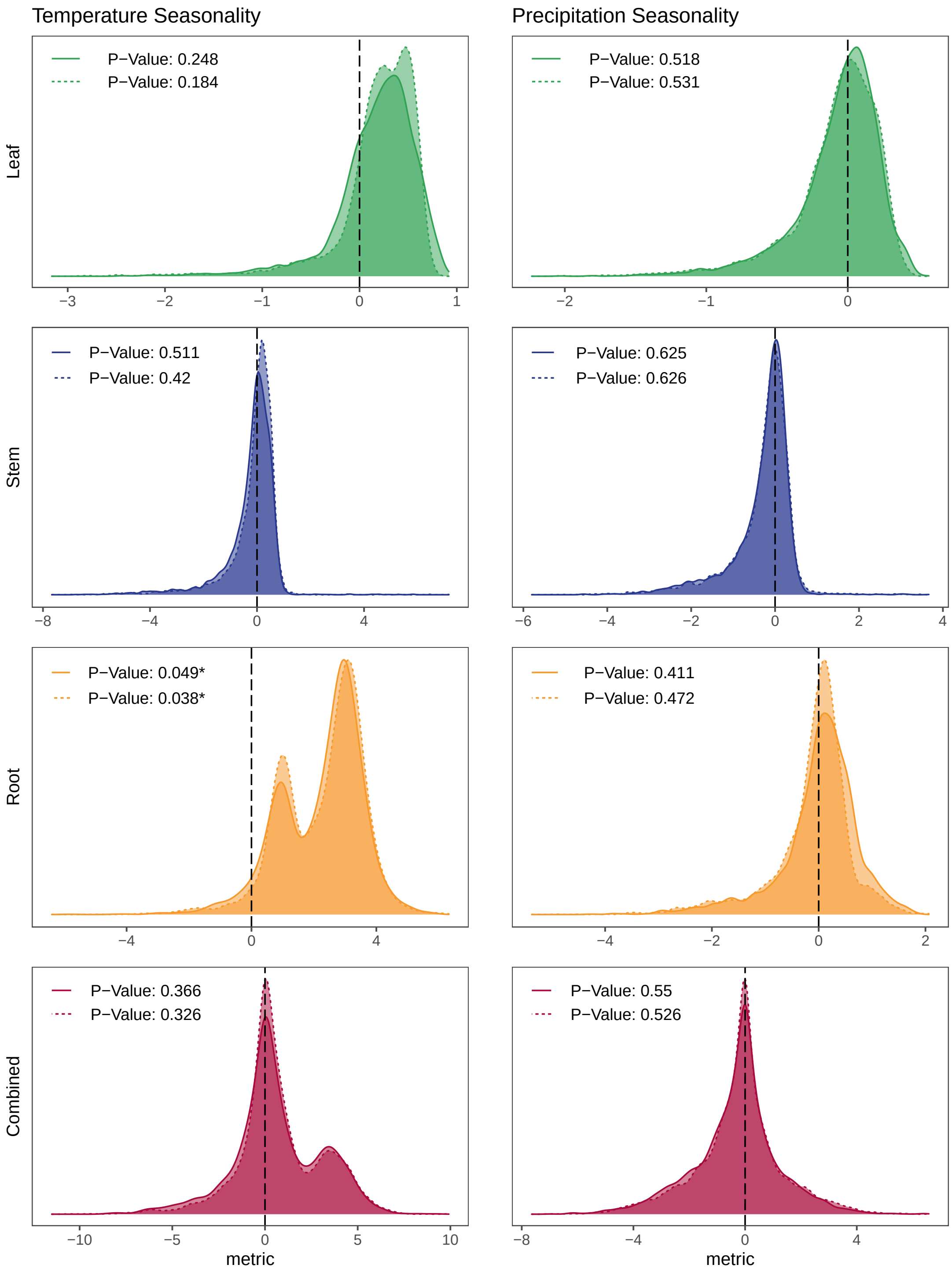


**Figure S2:** Test statistic ($S$) distributions for the first (solid lines) and second (dashed lines) sets of five phylogenies. Distributions with 95% $> 0$ are considered evidence for a statistically significant difference in state-specific $\theta$. P-Values correspond to the percent of values equal to or less than zero. These results are very similar to those presented in the main text. Root and temperature seasonality are significant while all other analyses are not significant.
